# Lymphocyte-Specific Biomarkers Associated with Preterm Birth and Bronchopulmonary Dysplasia

**DOI:** 10.1101/2020.11.03.367201

**Authors:** Soumyaroop Bhattacharya, Jared A. Mereness, Andrea M. Baran, Ravi S. Misra, Derick R. Peterson, Rita M. Ryan, Anne Marie Reynolds, Gloria S. Pryhuber, Thomas J. Mariani

## Abstract

Many premature babies who are born with neonatal respiratory distress syndrome (RDS) go on to develop Bronchopulmonary Dysplasia (BPD) and later Post-Prematurity Respiratory Disease (PRD) at one year corrected age, characterized by persistent or recurrent lower respiratory tract symptoms frequently related to inflammation and viral infection. Transcriptomic profiles were generated from sorted peripheral blood CD8+ T cells of preterm and full-term infants enrolled with consent in the NHLBI Prematurity and Respiratory Outcomes Program (PROP) at the University of Rochester and the University at Buffalo. We identified outcome-related gene expression patterns following standard methods to identify markers for oxygen utilization and BPD as outcomes in extremely premature infants. We further identified predictor gene sets for BPD based on transcriptomic data adjusted for gestational age at birth (GAB).

RNA-Seq analysis was completed for CD8+ T cells from 145 subjects. Among the subjects with highest risk for BPD (born at <29 weeks gestational age (GA); n=72), 501 genes were associated with oxygen utilization. In the same set of subjects, 571 genes were differentially expressed in subjects with a diagnosis of BPD and 105 genes were different in BPD subjects as defined by physiologic challenge. A set of 92 genes could predict BPD with a moderately high degree of accuracy. We consistently observed dysregulation of NRF2, HIPPO and CD40-associated pathways in BPD. Using gene expression data from both premature and full-term subjects (n=116), we identified a 28 gene set that predicted the PRD status with a moderately high level of accuracy.

Transcriptomic data from sort-purified peripheral blood CD8+ T cells from 145 preterm and full-term infants identified sets of molecular markers associated with independent development of BPD in extremely premature infants at high risk for the disease and of PRD among the preterm and full-term subjects.

## Introduction

Acute and chronic respiratory morbidities are common in extremely premature infants. (1) Increased survival of very premature infants is leading to increasing numbers of children with chronic lung disease. Since the end of the last millennium, the rate of premature births <34 weeks of gestation have consistently increased in the United States, and in 2008 it was 12.3%. (2) Among the extremely preterm infants, 20-35% die before their discharge home (3,4). Prematurity-related deaths accounted for 35% of all infant deaths in 2010, more than any other single cause. Preterm birth cost the U.S. health care system more than $26 billion in 2005 (5). Among NICU survivors, approximately 40% develop Bronchopulmonary Dysplasia (BPD), a chronic lung disease of the newborn. BPD has both genetic and environmental risk factors. It is characterized by varying degrees of lung injury potentially due to required supplemental oxygen and mechanical ventilation and is often associated with infection. BPD results from abnormal repair and impaired lung development after acute lung injury. Airway function may even deteriorate during the first year of life in infants with BPD (6). Infants with BPD are more likely to die than those without chronic lung disease, even if they survive the initial hospitalization.

The NIH NHLBI Prematurity and Respiratory Outcomes Program (PROP) enrolled 835 extremely premature infants and collected multiple biospecimens over time and extensive data including BPD and respiratory morbidity outcomes over the 1st year of life. PROP investigated the molecular mechanisms contributing to the risk of respiratory disease in premature neonates over the first year of life. A set of clinical and non-invasive respiratory assessments were performed, based on the respiratory status of the infant at the time of testing, and was used to predict the severity of respiratory outcomes in the first year of life. The primary goal of the PROP studies was to identify biomarkers (biochemical, physiological and genetic) that are associated with, and thus potentially predictive of, respiratory morbidity in preterm infants up to 1-year corrected age. A validated, objective measure of pulmonary outcome at 1 year does not currently exist. In addition to the identification of markers of BPD, one of the primary outcomes in the PROP study was defined as presence or absence of Post-Prematurity Respiratory Disease (PRD). In order to be classified as having PRD, infants were required to have a positive response in at least 1 of the 4 morbidity domains ((1) hospitalization for respiratory indication, (2) home respiratory support, (3) respiratory medication administration, and/or (4) respiratory symptoms without a cold) during at least 2 separate parental interviews conducted at 3-, 6-, 9- and 12-months corrected age (7).

High throughput sequencing for genome-wide transcriptomic analysis, by RNA-Seq or microarrays, is an unbiased approach applied to identify biomarkers that may provide predictive value. These approaches have been proven to be powerful tools capable of biomarker discovery for various disease states including BPD. We have previously presented an analysis of lung tissue gene expression in subjects with BPD (8). Application of blood-based gene expression profiling can potentially provide novel biomarkers for diagnosis and therapeutic management of BPD. Previous studies have used whole blood–derived peripheral blood mononuclear cells (PBMC) as a means of mining for novel markers for BPD (9). PBMCs are relatively easy to obtain from whole blood and can be sorted into leukocytes, including B cells, T cells, monocytes, and natural killer cells (10). Emerging data suggest an important role for T lymphocytes in the pathogenesis of chronic lung disease in babies born prematurely. CD8+ T cells, TNF-α, TNF receptors and NK cells provide protection from viral infection but also contribute to the immunopathology, by contact-dependent effector functions (perforin and FasL).

IFN-γ and particularly TNF-α are thought to be primary perpetrators of T-cell-mediated lung injury (11). CD8+ T cells have shown adaptive immune insufficiency in newborn mice infected with influenza A within one week of birth (12). RSV infected neonatal mice recruited CD8+ T cells defective in IFN-γ production in association with mild symptoms. Re-infection as adults however resulted in limited viral replication but enhanced inflammation and T cell recruitment, including Th2 cells and eosinophils (13,14). Depletion of CD8+ T cells (but not CD4) cells during the primary neonatal infection was protective against the adult challenge. We have previously shown that CD8+ T cells appear to play a pathogenic role in subjects with BPD, and may be associated with overall risk for lung morbidity (15). In related studies, we have observed that CD8+ T cells are increased in both mouse and human lungs exposed as neonates to hyperoxia, and have a hyper-responsive, fibrotic and destructive response to subsequent viral infection (8,16). Further, these cells have a predominant role in direct cytotoxicity in the lung, via interactions with epithelial cells and as regulators of macrophage responses, as well as in general human resistance to viral infection. In a separate population of premature infants, enrolled in the PROP study, phenotyped T cells at birth, at 36 weeks of adjusted gestational age, and at 12-months corrected age, were associated with a PROP-defined respiratory morbidity at 12 months (17). The goal for the current study was to demonstrate that transcriptional profiling of CD8+ T cells, obtained from premature infants at discharge, can identify disease-related gene expression patterns informative for pathogenesis and capable of predicting risk of future respiratory distress. We hypothesized that transcriptomic analysis of sorted lymphocyte sub-populations could identify predictive markers and pathways associated with respiratory outcomes. We followed a cohort of 157 infants, ranging from 23 to 41 weeks of gestation at birth, enrolled in the Prematurity and Respiratory Outcomes Program (PROP) at the University of Rochester Medical Center and Children’s Hospital of Buffalo (18). Here, we present a novel gene expression RNA-seq data set generated from CD8+ T cells from 145 subjects with varying levels of premature birth and report the identification of disease biomarkers for BPD and PRD.

## Methods

This study aimed at generating transcriptomic profiles of CD8+ T cells from newborn human blood. This study was approved by the Institutional Review Board of University of Rochester with a Memorandum of Understanding executed with the University of Buffalo IRB. Subjects were enrolled within gestational age at birth (GAB) epochs, in order to characterize the relationship among prematurity, disease risk and gene expression. The steps involved in the process, starting from sample collection to isolation of total RNA have been described in detail in our previous publication (10). Additional steps relevant to this study are shown in Supplemental Figure 1.

### Oxygen Exposure, BPD Diagnosis & PRD

The traditional categorical approach of classifying BPD as absent or present is likely an oversimplification. Tooley (19) recommended that oxygen use at 28 days of age would identify preterm infants with BPD. Almost a decade later, Shennan and colleagues proposed that the best predictor of abnormal pulmonary outcomes among very low birth weight premature infants was the clinical use of oxygen at 36 weeks postmenstrual age (PMA) (20). A workshop convened by the National Institutes of Health (NIH) proposed severity-based diagnostic criteria for BPD (21) that included the use of oxygen for at least 28 days (not necessarily consecutive) and an assessment of respiratory support at 36 weeks PMA, recognizing that some infants breathing room air at 36 weeks PMA may have residual lung disease. The majority of infants with birth weights less than 1 kg will have a diagnosis of at least mild BPD by the Consensus Conference definition based on 28 days in oxygen (21). Given clinical variations in oxygen administration, a structured trial of room air test was developed by the NICHD Neonatal Research Network (22), the frequency of BPD among the subjects was determined using two previously published definitions: the Shennan definition (20), which defines BPD as supplemental oxygen requirement at 36 weeks PMA in infants born with birth weight (BW) less than 1500 grams, and a physiologic definition with a room-air challenge (RAC) which defines BPD as requirement of oxygen support (>21% O_2_) for at least 28 days and a subsequent assessment at 36 weeks PMA or discharge, whichever comes first. (7,22),.

NICU oxygen exposure was calculated, as previously reported, from the FIO_2_ recorded in the medical record once each noon for the first 14 days of life. FIO_2_ was corrected to Effective FiO2 for low nasal cannula flow using established tables (23). Oxygen utilization or Oxygen_AUC_ was calculated by the formula defined in Benaron and Benitz (1994) (24) using information recorded in the daily respiratory flowsheet data (FIO2, respiratory support mode, and applied airway pressure or cannula flow) through the first 28 days of life. We chose to look at Oxygen_AUC_ at 14 days of postnatal age (Oxygen_AUC14_) to include the second postnatal week to capture pulmonary deterioration as presented in BPD (25).

The infants enrolled in the study were followed up periodically for up to 12 months of age, corrected by gestational age (CGA) at birth. At the 12-month CGA follow-up visit they were assessed for persistent respiratory distress based on the frequency of hospitalization due to any kind of respiratory distress. Persistent Respiratory Distress (PRD) was diagnosed if there were positive responses in at least one of the following domains: (1) hospitalization for respiratory indication, (2) respiratory support at home, (3) respiratory medication administration, and/or (4) cough or wheeze without a cold, reported on at least two caregiver post-discharge surveys completed at 3, 6, 9 and 12 months CGA, as previously reported (7). The definitions and criteria for the different diagnoses have been listed in Supplemental Table 1.

### Sample Collection and RNA Isolation

An average of 2.5 ml of venous blood was collected into sodium heparin glass vacutainers from premature infants enrolled with consent in the Prematurity and Respiratory Outcomes Program (PROP), at the time of hospital discharge at the University of Rochester and the University at Buffalo and shipped to a central laboratory in Rochester. Freshly purified PBMCs were isolated by Ficoll gradient (Amersham Pharmacia Biotech # 17-1440-03) centrifugation, from the whole blood diluted 1:2 with 1x dPBS, and counted according to previously established protocols (26). In subjects with at least 8 million cells, 5 million cells were stained with antibodies to individual lymphocyte markers, and sorted on a FACSAriaII sorter at the Flow Cytometry Core facility of the University of Rochester as previously reported (10). CD3+CD8+, CD3+CD4+, CD3-CD56+ (NK) or CD3-CD19+ (B) cells were collected separately. Non-marker positive and dead cells were discarded. Sorted cells were spun into pellets, which were further lysed and frozen. The steps from collection to lysis of each sample were completed within a 24-hour period in order to preserve RNA quality and integrity. Frozen lysates were thawed and RNA was extracted using Agilent Absolute RNA Microprep kit (catalog #400805), with an on-column DNase digestion, as per manufacturer recommended protocol.

### RNA-Seq and Data Generation

For the current study, RNA isolated from the sorted CD8+ T cells from 145 pre-term and full-term subjects, was used for transcriptomic profiling by RNA-seq. cDNA libraries were generated with 1 ng RNA, using the SMARter Ultra Low Amplification kit (Clonetech, Mountain, CA). cDNA quantity was determined with the Qubit Flourometer (Life Technologies, Grand Island, NY) and quality was assessed using the Agilent Bioanalyzer 2100 (Santa Clara, CA). Libraries were sequenced (single endreads) on the Illumina HiSeq2500 (Illumina, San Diego, CA) to generate 20 million reads/sample. Reads generated from the sequencer were aligned using the TopHat algorithm (27) and expression values were summarized using HTSeq (28). The data from this study has been provided in dbGAP. The dbGaP accession assigned to this study is phs001297.v1.p1.

### Normalization and Filtering

Samples were excluded based on poor read count/mapped read numbers, or if they were extreme outliers in hierarchical clustering and Principal Components Analysis (PCA). Genes were excluded if they were not consistently identified as expressed (a count of zero in over 1/3 of subjects). Subjects with high prevalence (>75%) of zero/low reads (raw count value ≤5) were excluded. Genes with high prevalence (>75%) of low counts (normalized count value ≤3) across subjects were excluded.

The subject-specific conditional upper quartile (UQ, 75^th^ percentile) among non-zero reads was computed. The subject-specific normalization factor was calculated by dividing the UQ for a given subject by the mean UQ across all subjects. The normalized gene values for a given subject were calculated by dividing the raw count values by the normalization factor for that subject. After adding 1 to all values to account for zeros, the normalized counts were log_2_ transformed.

### Selection of Univariately Differentially Expressed Genes

Differences in gene expression between subject groups was assessed by SAM-Seq (29) and Likelihood Ratio Test (LRT). SAM-Seq was used to identify genes with expression patterns significantly (FDR<0.05) associated with BPD, RAC. LRT for log (normalized RNA-Seq), adjusted for GAB via logistic regression, was used to identify genes with expression patterns significantly (FDR<0.05) associated with BPD in subjects born at less than 29 weeks of age. For quantitative analysis, the correlations between normalized counts and oxygen utilization at 14 days were estimated.

Expression changes of the genes, identified as significantly different in BPD by the tests, were assessed in transcriptomic profiles of PBMCs obtained from infants with BPD, and age matched controls generated from an independent cohort (9), using the data available on Gene Expression Omnibus (GSE32472).

### Prediction of Bronchopulmonary Dysplasia (BPD) via Screened Principal Components

The following method is our minor variant of *Screened Principal Components Analysis (sPCA)* (30), where the genes were univariately screened, and those with a nominal Wilcoxon p□<□0.10 were used for further analysis. The first principal component (PC1) of the genes was derived, and genes with loadings close to 0 were removed. Genes most strongly univariately associated with BPD (with or without adjusting for gestational age) were selected using a screening threshold chosen by cross-validation (CV). The first Principal Component (PC1) of the genes passing the univariate screen was constructed and a logistic regression model was fit to predict BPD as a function of PC1 (and optionally gestational age). Receiver Operating Characteristic (ROC) curves depicting sensitivity and specificity along with associated AUC were estimated without (naïve AUC) and with an outer loop of nested CV (CV-AUC).

### Prediction of Post-Prematurity Respiratory Disease (PRD) via Canonical Pathways

We have used canonical pathway analysis where we grouped our 13,434 genes into 1,330 biologically-relevant pathway-based gene sets of molecular signature database (mSigDB) (31), where each gene belongs to 0, 1, or more pathways. We then reduced the constituent genes in each pathway to their PC1. Genes belonging to 0 pathways were thus excluded from consideration, while CV screened logistic forward selection (or LASSO) was applied to the 1330 pathway-based PC’s. Bivariate CV was used to simultaneously select both the threshold for univariate logistic likelihood ratio test screening of pathways (with or without adjusting for gestational age) and the final number of pathways chosen by forward selection (or the LASSO penalty). The entire procedure was then nested within an outer loop of nested CV in order to estimate performance via the Receiver Operating Characteristic (ROC) curve and its associated Area Under the ROC Curve (CV-AUC). This method has been described in details in our previous publication (21).

### Functional Classification

Genes identified as differentially expressed in individual comparisons were used for independent functional classification through canonical pathway analysis, and upstream regulators identification using Ingenuity Pathway Analysis (IPA; Qiagen, Carlsbad, CA). Pathways and upstream regulators were identified as significantly associated with the diagnoses by Fisher’s exact test (p < 0.05 or -log(p-value) > 1.3) as calculated by IPA.

### Quantitative reverse transcriptase–polymerase chain reaction (qPCR)

cDNA was synthesized from 100 ng RNA using iScript cDNA synthesis kit (Biorad, HerculesCA) and quantitative reverse transcriptase–polymerase chain reaction (qPCR) was performed with a Viia7 (Applied Biosytems, Carlsbad, CA) using SYBR green chemistry as previously described (8) using noncommercial (http://pga.mgh.harvard.edu/primerbank) assays. Gene expression levels were calculated relative to GAPDH as an internal, endogenous control, according to the ddCT method.

## Results

### Subject Demographics

Peripheral blood was collected at the time of first hospital discharge, from 145 preterm and full-term infants enrolled in the NHLBI Prematurity and Respiratory Outcomes Program (PROP) at the University of Rochester and the University at Buffalo. Of the 145 subjects, 72 were extremely premature having been born at less 29 weeks of gestation. There was insufficient evidence that the distribution of race (p=0.64) or sex (p=1) differed between BPD and non-BPD subjects, while as expected gestational age at birth was lower for those with BPD (p=0.03). Similarly, among all subjects (n=130) there was insufficient evidence of any difference in race (p=0.70) or sex (p=1) by PRD status, while as expected gestational age at birth was lower for those with PRD (p=0.01; Table 1). Detailed diagnostic and demographic information for the individual subjects in provided in Supplemental Table 2.

**Table 1:** Subject Demographics

| **Subjects with**<br>**GAB < 29 Weeks (n=72)** | **BPD**<br>**N=34** | **Non BPD**<br>**N=38** | **P value\*** |
| --- | --- | --- | --- |
| GAB in Weeks, mean ± SD<sup>b</sup> | 26.1 ± 1.4 | 26.9 ± 1.5 | 0.03 |
| Male Sex, No. (%) | 17 (50) | 18 (47) | 1.00 |
| White Race No. (%) | 20 (59) | 20 (53) | 0.64 |

| **Demographics**<br>**All Subjects (n=130)** | **PRD YES**<br>**N=70** | **PRD NO**<br>**N=46** | **PRD NA<sup>#</sup>**<br>**N=14** | **P value\*** |
| --- | --- | --- | --- | --- |
| GAB in Weeks, mean ± SD | 28.5 ± 3.3 | 31.1 ± 5.2 | 32.2 ± 4.1 | 0.01 |
| Male Sex, No. (%) | 34 (48) | 23 (50) | 10 (71) | 1.00 |
| White Race, No. (%) | 43 (61) | 30 (65) | 9 (64) | 0.7 |

\* Fisher's Exact or t-test P-value
GAB: gestational age at birth
PRD NA<sup>#</sup>: PRD Status Not Available

Transcriptomic profiles were generated from sorted and purified CD8+ T cells obtained from the blood collected at discharge. The analytical data set includes values from 13,455 genes for 130 samples, post filtering. As reported previously, the average number of sequence reads in the samples were high (9.93±3.69 million sequence reads). Overall, approximately 60% of possible genes showed detectable transcript as expected for a subset of differentiated cell type (10) as shown in Supplemental Figure 2.

### Molecular Markers for BPD

Gene expression patterns associated with cumulative oxygen exposure (over the first 14 days of life) in subjects at greatest risk for BPD (born at GAB<29 weeks; n=72) was assessed. Rank correlation analysis identified 501 genes to be significantly associated (at pFDR<0.1) with oxygen exposure, of which 403 were upregulated in BPD, while 98 genes were downregulated in BPD. The magnitude of change, however, was not large, with only 1 gene (*GPCPD1*) induced by 2.3 fold, while all other changes were less than 2-fold, irrespective of the directionality. Twelve of these genes (*RETN, EPHX2, CD27, NOSIP, APOA1BP, TMCO6, KLHL3, B3GALNT1, SLC9A4, PRKCD, ZNF791, and B3GNT2*) were also identified as differentially expressed in BPD in an independent study looking at BPD markers in PBMCs (9). The 501 genes, when further assessed for functional classification by Ingenuity Pathway Analysis (IPA), were found to be associated with 104 pathways and 300 upstream regulators (Figure 1). The pathways associated with oxygen utilization included epithelial-mesenchymal transition, and multiple immune signaling pathways, while tubule formation associated and immunologic molecules were present among the upstream regulators.

**Figure 1:**
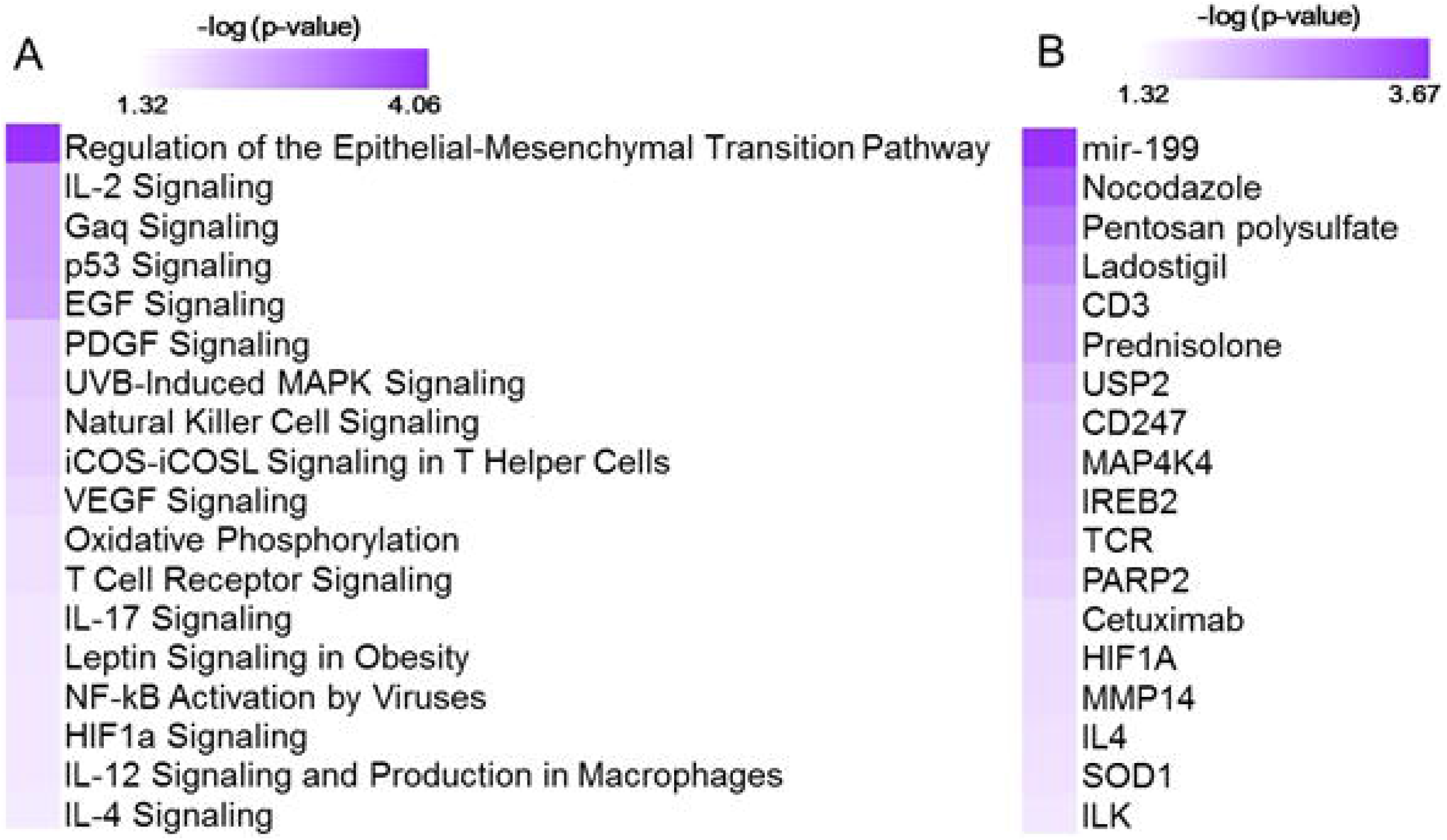
Functional analysis using 501 genes associated with oxygen utilization at day 14 (Oxygen_AUC14_) identified with 104 pathways and 300 upstream regulators. Shown here are selected significant canonical pathways (A) and upstream regulators (B), along with their significance level (-logP) as generated by IPA.

Data from 72 subjects born at <29 weeks CGA was used to identify gene expression associated with BPD as defined by physiologic challenge (RAC) or by Shennan criteria. Using SAM-Seq (at mFDR<0.1), 571 genes were differentially expressed in subjects receiving a diagnosis of BPD (Shennan). While all of the 571 genes were upregulated in BPD, only five genes (*GPCPD1, TMEM2, USP2, TSPYL2, and ELL2*) showed a magnitude of induction of greater than 2.0 fold. Fourteen of these genes (*RNF125, FEM1C, FAM54A, ZNF791, AXIN2, B3GNT2, ZNF565, SPON1, TIPARP, ZBTB3, FAM115C, PELO, MXD1, and PFKFB3*) were also identified as differentially expressed, and over expressed in BPD in an independent study looking at BPD markers in PBMCs (9). These 571 genes, when further assessed for functional classification by IPA, identified 113 canonical pathways and 409 upstream regulators to be associated with BPD.

In addition, 105 genes were differentially expressed (SAM-Seq at mFDR<0.1) in subjects who failed a room air challenge. While all of the 105 genes were upregulated in BPD, only two genes (*GPCPD1, and TSPYL2*) showed a magnitude of induction of greater than 2.0 fold. These 105 genes when analyzed by IPA, identified 18 canonical pathways and 415 upstream regulators (Figure 2). Among the pathways associated with BPD, PEDF, CD40, PI3K/AKT, VEGF and NF-kB signaling were predicted to be activated, while p53 signaling was inhibited in BPD. Among the BPD associated upstream regulators, CD23, CD28, PTEN, and TCR, are inhibited, while NFκB inhibitor, camptothecin, and dexamethasone were activated in BPD. Activation of camptothecin indicates potential inhibition of topoisomerase I, a molecule playing critical role in development, and maintenance of transcription from supercoiled DNA. This may potentially explain the developmental arrest in lungs from subjects with BPD. Activation of dexamethasone may be an indicator of stress response due to steroid application as therapeutic response to BPD. When adjusted for gestational age at birth, 75 genes were associated with the diagnosis of BPD (Shennan) as identified by the Likelihood Ratio Test (LRT at FDR<0.1), and were upregulated in BPD. On further analysis by IPA, the genes provided 113 canonical pathways and 409 upstream regulators (Figure 3), of which neuroinflammation pathway appeared to be inhibited, while upstream regulators, CD23, CD28, and TCR, are inhibited, and camptothecin, and dexamethasone were activated in BPD. Screened Principal Components Analysis (sPCA), with screening adjusted for GAB, outperformed both screened LASSO and forward selection and identified a classifier gene set consisting of 92 genes (naïve AUC=0.86; CV-AUC=0.71) associated with BPD (Shennan), representing 21 canonical pathways and 253 upstream regulators (Supplemental Figure 3). All of the 92 genes were upregulated in BPD, however, none of the genes except *GPCPD1* had a magnitude of induction of greater than 2fold. These 92 genes were also inclusive of all the 75 genes identified by LRT.

**Figure 2:**
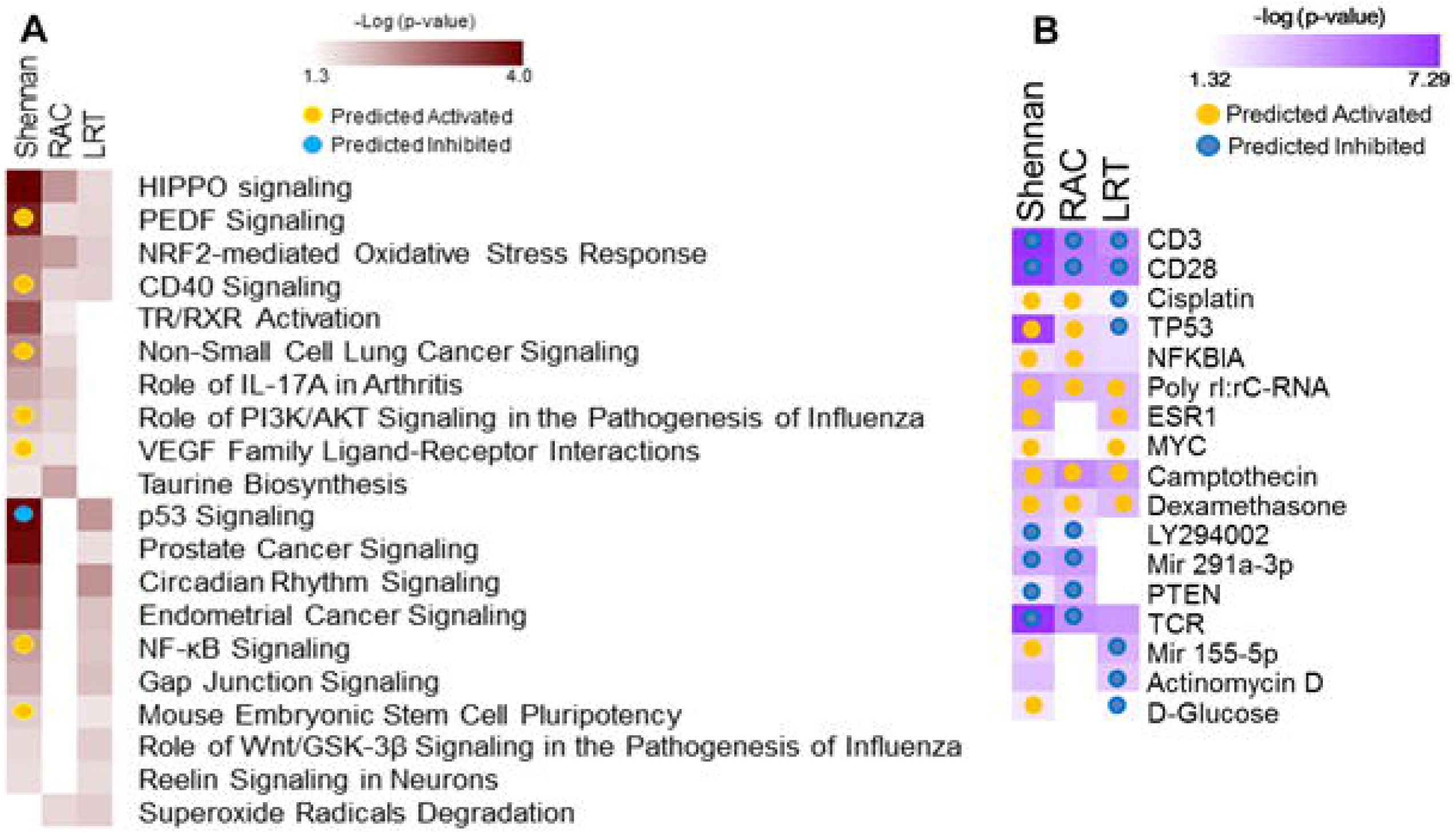
Functional analysis of genes associated with BPD. Separate analyses were performed for gene sets identified from analysis of 571 genes associated with BPD (Shennan) and 101 genes associated with RAC as identified by SAM-Seq, and 92 genes identified by sPCA (adjusted for GAB) to be associated with BPD (Shennan). Selected canonical pathways (A) and upstream regulators (B) identified are listed for each analysis (columns), with significance (-logP) and directionality (activated/inhibited) as generated by IPA.

**Figure 3:**
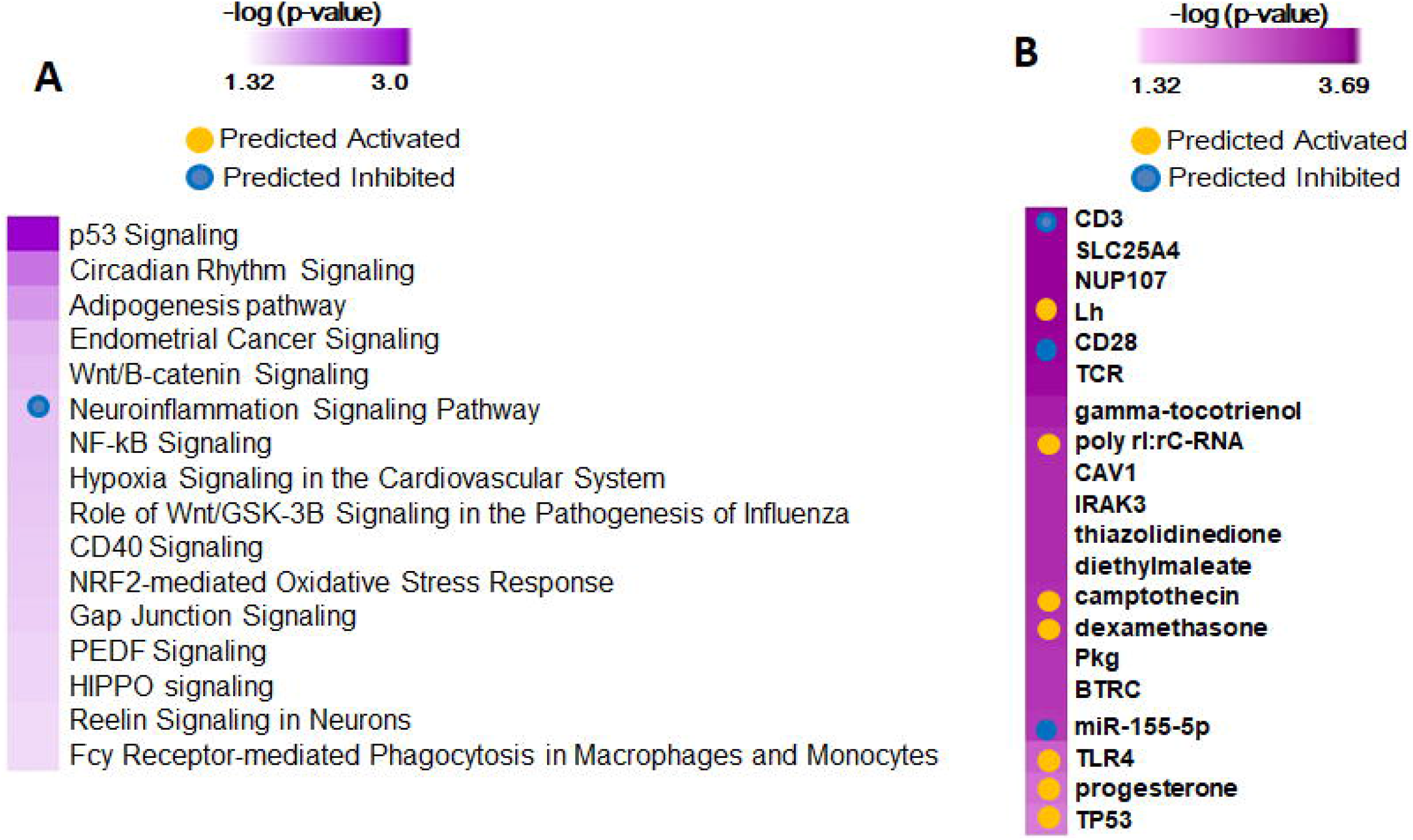
Functional analysis of 75 predictor genes for BPD defined by LRT, identified 113 canonical pathways and 409 upstream regulators. Selected significant canonical pathways (A) and upstream regulators (B) identified are shown, along with their significance level (-logP) as generated by IPA.

We subsequently assessed gene expression changes in BPD based on multiple physiologic and clinical definitions and were successful in identifying nine genes (*GPCPD1, MTSS1L, USP15, DDX24, KLF9, CLK1, ZC3H7A, ITCH, and PIK3R1*) that were consistently different, and upregulated in BPD, irrespective of definitions, or analytical approaches (Supplemental Figure 4).

### Classifiers for PRD Status

Gene expression data from all subjects with PRD status (n=116), was used to identify a set of marker genes, based on our novel *canonical pathway analysis* (32), in order to classify the subjects by PRD status (PRD: n=70 and No PRD: n=46). Screened logistic forward selection outperformed both screened LASSO and sPCA, and gestational age was excluded since it did not improve performance. This process identified a set of 28 genes, derived from four canonical pathways (Table 2), which predicted PRD status with a moderately high degree of accuracy (naïve AUC=0.85; CV-AUC=0.70) (Figure 4). Interestingly, gene predictors of PRD were associated with pathways involving complement activation, organic ion transport and regulation of intracellular actin organization.

**Table 2:**
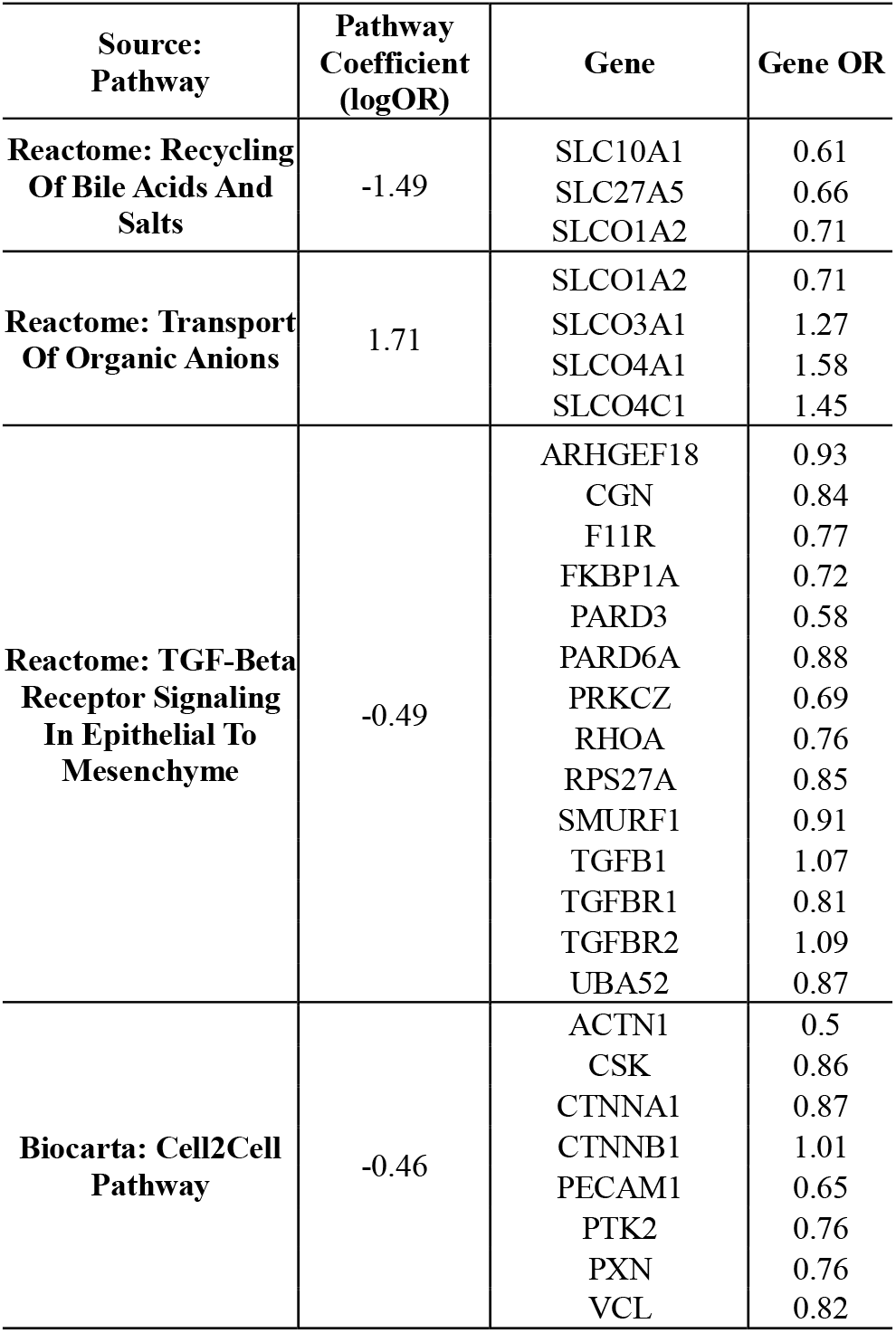
Our novel *canonical pathway analysis* selected 4 pathways including a total of 28 genes using cross-validated screened logistic forward selection for prediction of PRD Status. Shown are pathway and gene names, estimated pathway coefficients (logOR), and constrained gene odds ratios (OR) factoring in the PCA loadings for each gene within a pathway.

**Figure 4:**
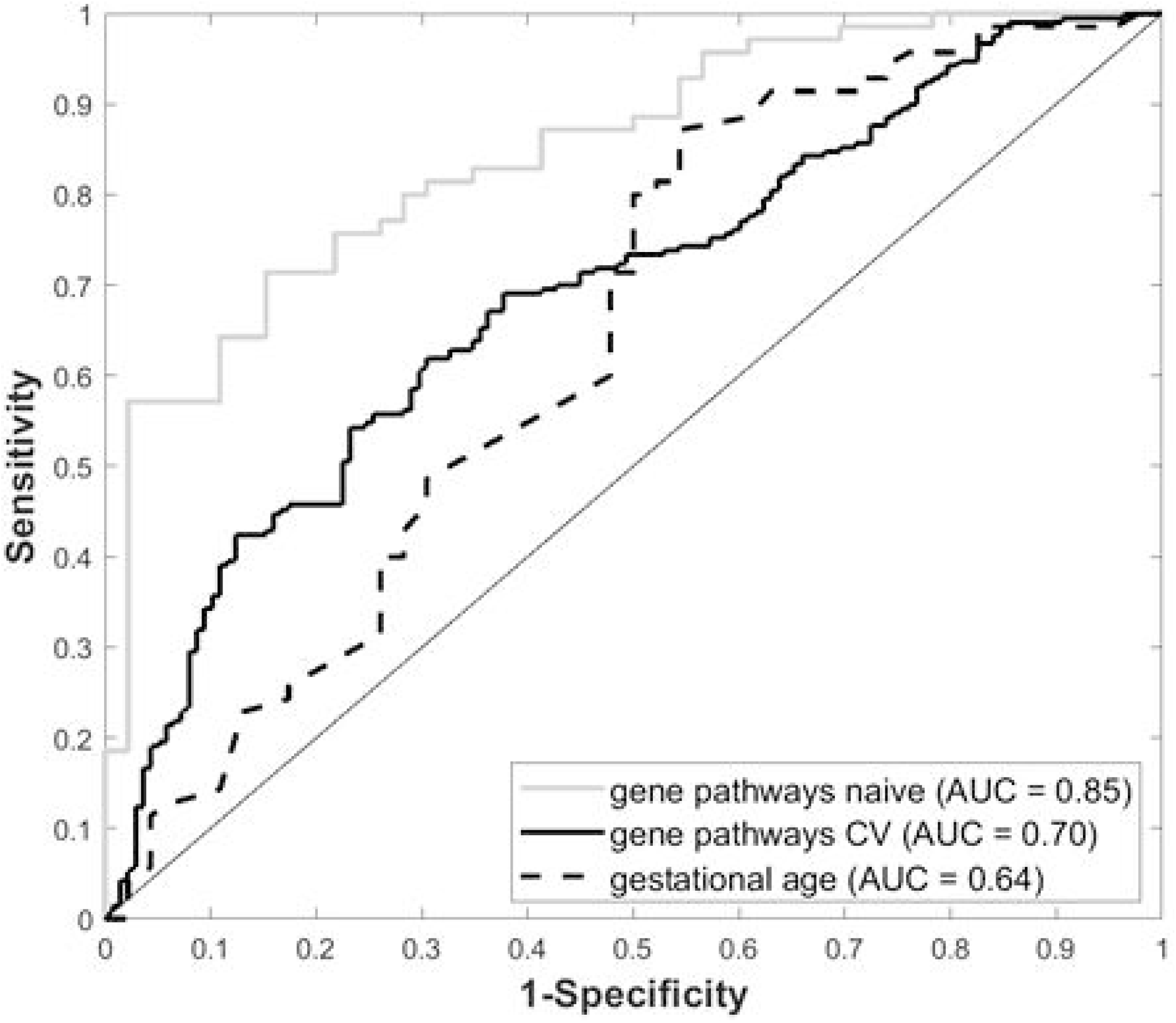
Receiver Operating Characteristic (ROC) curves for gestational age and our 4-pathway 28-gene model, with associated Area Under the ROC Curve (AUC).

### qPCR Validation

Molecular validation of the BPD-associated transcriptomic changes was attempted on a set of eleven genes selected based on their magnitude of difference or biological relevance, by quantitative reverse transcriptase-polymerase chain reaction (qPCR) (Table 3). For each gene, the UQ normalized RNA-seq counts were correlated with the gene expression levels determined by qPCR of the CD8+ T cell cDNA as defined by the dCt. GAPDH was used as the endogenous control or housekeeping gene, whose Ct was subtracted from gene Ct to determine the dCT values for each of the genes tested. Spearman’s rank correlation coefficient was estimated along with an associated p-value for each gene. We observed validation of the sequence data for 9 of the 11 genes, as defined by a significant Spearman rank correlation (p<0.05) in expression between sequence-based and qPCR-based expression levels. Three genes (*KLF9, DLG5 and ZNF44*) differed in expression between BPD and non-BPD subjects (p<0.05), while one additional gene (*PSME4*) had borderline yet insufficient evidence of a difference (0.05<p<0.10).

**Table 3:**
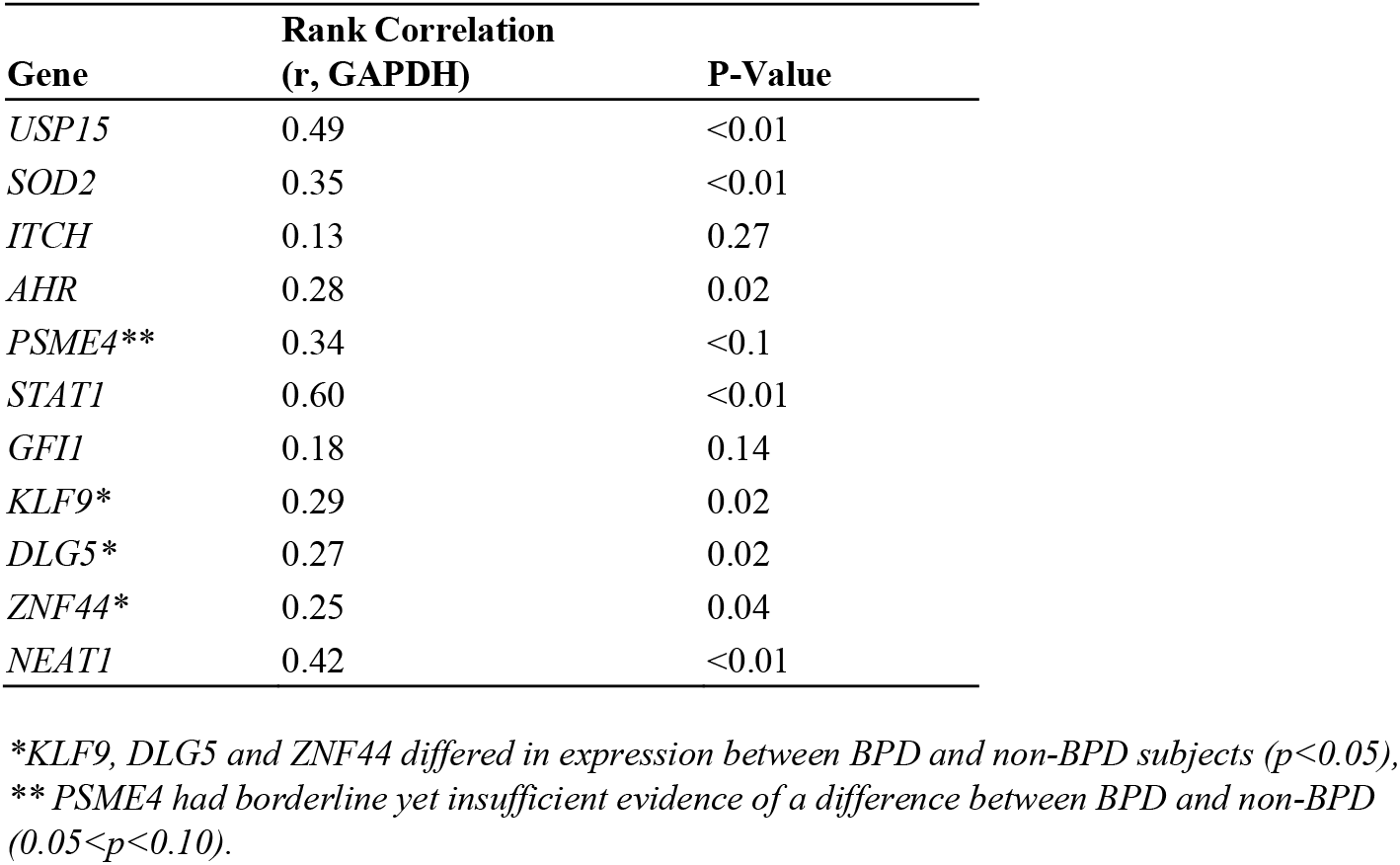
Validation of BPD Markers

## Discussion

Premature birth is defined as birth taking place prior to 37 weeks of GAB. Prematurity associated lung diseases have been reported to affect not only children as newborns but to also predispose to prolonged respiratory morbidity later in life. Unfortunately, not much is known about the pathophysiology of the prematurity associated lung diseases, such as BPD, and other chronic and prolonged childhood respiratory diseases. Transcriptomic assessment using gene-expression microarrays has previously been used to identify markers for normal lung development as well as BPD (8,33,34). As an alternative to lung tissues, gene expression analyses using peripheral blood have been used in lung diseases to study pathogenesis, severity and recently as a diagnostic tool (32,35). In this study we have used high-throughput sequencing to explore peripheral gene-expression changes associated with prematurity. Our analysis identified 571 genes differentially expressed in subjects with diagnosed instances of BPD when compared to extremely low birth weight (ELBW) controls born at less than 29 weeks of GAB by SAM-Seq. The list is further restricted to 92, when we adjust for gestational age at birth (sPCA) as shown in Supplemental Figure 4. In addition, we have identified markers, pathways and upstream regulators putatively associated with cumulative oxygen utilization.

Among the differentially expressed genes associated with BPD, *PFKFB3* gene was not only consistently identified as differentially expressed in BPD subjects irrespective of the approach used, but was also identified as a BPD marker in an independent transcriptomic analysis of PBMCs derived from BPD subjects (9). Through murine studies *PFKFB3* has been identified as potential therapeutic target for the treatment of Pulmonary Hypertension (PH) (36) Increased expression PFKFB3 in BPD is consistent with PH associated with BPD which is characterized by abnormal vascular remodeling, and vascular growth arrest, which are well documented pathophysiology associated with BPD (37). One of the differentially expressed genes, *KLF9*, has been previously identified as differentially expressed in T cells from patients with autoimmune rheumatoid arthritis (38), while another, DLG5, is involved in the HIPPO pathway. In fact, pathway analyses indicate dysregulation of *NRF2, HIPPO and CD40* pathways to be consistently associated with BPD. Another marker, *PSEM4*, has been associated with tuberculosis through *in vitro* and *in vivo* cultures, and is also related to CD4, IFNbeta1, and TGFbetaI pathways (39). The pathway most associated with oxygen utilization is the epithelial-mesenchymal transition pathway, which is unexpected, considering we are measuring gene expression in immune cells, but may potentially indicate underlying association with TGF or integrin pathways. While IL2 signaling is expected, Gaq is known to play a role in survival of immune cells (B and T-cells) (40,41). In addition, iCOS-iCOSL are known to be involved in T-cell skewing, and reduced SOD expression is related to impaired CD8+ T cell responses in tumor infiltrating lymphocytes (42). Interestingly, regulators associated by anti-survival, dexamethasone and camptothecin, appear to be activated in BPD, while T cell co-receptors, CD3 and CD28, appear to be inhibited in BPD. These may indicate potential arrest in lung development as a consequence of BPD and indicate induced immune and stress response as a result of therapeutic responses to BPD, due to either oxidative stress, or surfactant treatment.

We explored multiple approaches to identify a set of genes whose expression may be useful for classification of markers associated with prolonged respiratory morbidity in infants born prematurely. This included approaches to leverage biological priors as a means of identifying the most robust predictors since (1) expression changes at the individual gene level alone may not be sufficient to identify biologically meaningful data and (2) there exists substantial statistical advantages to dimension reduction strategies in the analysis of genome-wide data. We used a curated list of genes (43) to partition our transcriptomic data into 1330 biologically-relevant gene sets. Using a novel data reduction approach, we identified a 28 gene set classifier that groups subjects according to their PRD status with a moderately high degree of accuracy. The genes involved in the classifier gene sets have been known be associated with complement activation, organic ion transport, and regulation of intracellular actin organization, among others. Complement pathway components have long been known to be associated with immune responses leading to lung disorders, particularly asthma (44). Their association with persistent respiratory disease indicates propensity to future immunological complications, which may result in chronic lung disorders such as asthma. It has been reported that maintenance of normal ion transport in the lung epithelium also involves formation of thin layer coatings on the conducting airways (45). Indication of altered ion transport mechanism in respiratory disease, primarily due to prematurity, may lead to lung pathologies in future, in these subjects. Disruption of cellular organization during the embryonic development and childhood have long been associated with the origins of some adult lung diseases (46). Even with the limited set of predictive markers, it does appear that gene expression changes, in peripheral blood at the time of initial discharge after birth, are indicative of future respiratory diseases later in childhood and in adulthood.

One of the limitations of this study is the use of CD8+ T-cells in identifying biomarkers of a lung disease. While peripheral markers have been widely used in identifying expression based markers, it needs to be acknowledged that these hematopoietic cells are from different cell lineage from the pulmonary system, which is the primary organ system affected in BPD and subsequently in PRD. However, even with only one cell type of a different lineage, we have been able to identify previously known, as well as novel molecular markers, and pathways, associated with pulmonary disease due to premature birth. Another limitation of the study is the usage of multiple definitions of BPD in order to assess transcriptomic changes caused by it. Despite the various definitions and analytical approaches used, we were able to identify a set of nine marker genes that were observed to be increased in BPD, irrespective of the diagnostic definition and approach used. Through this study we have established proof-of-principal that gene expression provides value for predicting respiratory morbidity following pre-term birth.

In conclusion, we have successfully generated genome-wide transcriptomic data from sort-purified peripheral CD8+ T cells obtained from early pre-term, late preterm and term infants. We have identified molecular markers, pathways and upstream regulators putatively associated with cumulative oxygen utilization, BPD diagnosis, and PRD prediction. Further studies are needed to determine if the findings are unique to the circulating T cells sampled in this study, or reflective of similar effects in other cells including lung parenchymal cells.

## Author contributions

SB, JAM, AMB, and DRP analyzed the data. SB, RSM, and DRP wrote the paper. RMR, AMR, GSP and TJM conceived and designed the study. RSM, TJM and GSP oversaw the project.

## Acknowledgements

University of Rochester: Chin-Yi Chu, Heidie Huyck, Shelley Secor-Socha, Ernest Wang, Elizabeth Werner-Carbonell, Tanya Scalise, Dee Moffatt, Valerie Lunger, Sara Misra, Ashley Lopez, Jason Emo, Clement Ren (now at Indiana University), Carl D’Angio, William Maniscalco, Timothy Stevens, Sally Quataert, Kristin Scheible, Jeanne Holden-Wiltse, Sanjukta Bandopadhyay, the UR BLIS Data Management Team, UR RedCAP Database Team, UR Flow Core, UR Data Entry Team, UR Genomics Research Center

University at Buffalo: Jack Sharp, Shannon Castiglione, Mike Sacilowski, Aimee Horan, Karen Wynn, Patrick Conway, UB Data Entry Team

Funding: The Prematurity and Respiratory Outcomes Program (PROP) was supported by National Institutes of Health; National Heart, Lung, and Blood Institute (NHLBI); and *Eunice Kennedy Shriver* National Institute of Child Health and Human Development (U01 HL101794 [to University of Pennsylvania, B. Schmidt]; U01 HL101456 [to Vanderbilt University, J.L. Aschner]; U01 HL101798 [to University of California San Francisco, P.L. Ballard and R.L. Keller]; U01 HL101813 [to University of Rochester and University at Buffalo, G.P., R. Ryan, and T. Mariani]; U01 HL101465 [to Washington University, A. Hamvas and T. Ferkol]; U01

HL101800 [to Cincinnati Children’s Hospital Medical Center, A.H. Jobe and C.A. Chougnet]; and 5R01HL105702 [to Indiana University and Duke University, C.M. Cotton, S.D. Davis, and J.A. Voynow]). This research was conducted through cooperative agreements with NHLBI and in collaboration with the PROP Steering Committee. *The project described in this publication was supported by the University of Rochester CTSA award UL1 TR002001 from the National Center for Advancing Translational Sciences of the National Institutes of Health. The content is solely the responsibility of the authors and does not necessarily represent the official views of the National Institutes of Health*.

## PROP Investigators

Cincinnati Children’s Hospital Medical Center:

Barbara Alexander, RN, Claire Chougnet, PhD, Tari Gratton, PA, James M. Greenberg, MD, Cathy Grisby, BSN, CCRC, William Hardie, MD, Alan H. Jobe MD, PhD, Beth Koch, BHS, RRT, RPFT, Karen McDowell, MD, Kelly Thornton BS

Washington University:

Pamela Bates, CRT, RPFT, RPSGT, Claudia Cleveland, RRT, Thomas Ferkol, MD, Aaron Hamvas, MD, Julie Hoffmann, RN, Mark R. Holland, PhD, James Kemp, MD, Philip T. Levy, MD, Laura Linneman, RN, Jayne Sicard-Su, RN, Gina Simpson, RRT, CPFT, Gautam K. Singh, MD, Barbara Warner, MD

University of California at San Francisco:

Investigators

Philip L. Ballard, MD, PhD^1^, Roberta A. Ballard, MD^1^, David J. Durand, MD^2^, Eric C. Eichenwald, MD^4^, Roberta L. Keller, MD^1^, Amir M. Khan, MD^4^, Leslie Lusk, MD^1^, Jeffrey D. Merrill, MD^3^, Dennis W. Nielson, MD, PhD^1^, Elizabeth E. Rogers, MD^1^

Research Staff

Jeanette M. Asselin, MS, RRT-NPS^2^, Samantha Balan1, Katrina Burson, RN, BSN^4^, Cheryl Chapin^1^, Erna Josiah-Davis, RN, NP^3^, Carmen Garcia, RN, CCRP^4^, Hart Horneman^1^, Rick Hinojosa, BSRT, RRT, CPFT-NPS^4^, Christopher Johnson, MBA, RRT^4^, Susan Kelley, RRT^1^, Karin L. Knowles^1^, M. Layne Lillie, RN, BSN^4^, Karen Martin, RN^4^, Sarah Martin, RN, BSN^1^, Julie Arldt-McAlister, RN, BSN4, Georgia E. McDavid, RN4, Lori Pacello, RCP2, Shawna Rodgers, RN, BSN^4^, Daniel K. Sperry, RN4,^1^

^1^Department of Pediatrics, University of California San Francisco, San Francisco, CA;

^2^Children’s Hospital and Research Center Oakland, Oakland, CA

^3^Alta Bates Summit Medical Center, Berkeley, CA

^4^University of Texas Health Science Center-Houston, Houston, TX

Vanderbilt University:

Judy Aschner, MD, Amy B Beller BSN, Candice Fike, MD, Scott Guthrie, MD, Tina Hartert, MD, Nathalie Maitre, MD, Paul Moore, MD, Mark O’ Hunt, Theresa J. Rogers, RN, Odessa L. Settles, RN, MSN, CM, Steven Steele, RN, Marshall Summar, MD, Sharon Wadley, BSN, RN, CLS

University of Rochester-University at Buffalo:

Investigators:

Carl D’Angio, MD, Vasanth Kumar, MD, Tom Mariani, PhD, Gloria Pryhuber, MD, Clement Ren, MD, Anne Marie Reynolds, MD, MPH, Rita M. Ryan, MD*, Kristin Scheible, MD, Timothy Stevens, MD, MPH

Technical Staff:

Heidie Huyck, BS, Valerie Lunger, MS

Study Staff:

Shannon Castiglione, RN, Aimee Horan, LPN, Deanna Maffet, RN, Jane O’Donnell, PNP, Michael Sacilowski, MAT, Tanya Scalise, RN, BSN, Elizabeth Werner, MPH, Jason Zayac, BS

Respiratory Therapists and Nurses:

Kim Bordeaux, RRT, Pam Brown, RRT, Julia Epping, AAS, RT, Lisa Flattery-Walsh, RRT, Donna Germuga, RRT, CPFT, Nancy Jenks, RN, Mary Platt, RN, Eileen Popplewell, RRT, Sandra Prentice, CRT

*Present address, Case Western Reserve University, Cleveland, OH

Duke University:

Kim Ciccio, RN, C. Michael Cotten, M.D., Kim Fisher, Ph.D., Jack Sharp, M.D., Judith A. Voynow, M.D.*

*Present address, Virginia Commonwealth University

Indiana University:

Charles Clem, RRT, Stephanie Davis, M.D., Susan Gunn, NNP, CCRC, Lauren Jewett, RN, CCRC Brenda Poindexter, M.D., M.S.#

#Present address, University of Cincinnati

Steering Committee Chair:

Lynn M. Taussig, MD, University of Denver

NHLBI Program Officer:

Carol J. Blaisdell, MD

University of Pennsylvania Data Coordinating Center:

Scarlett Bellamy, ScD, Maria Blanco, BS, Denise Cifelli, MS, Sara DeMauro, MD,

Jonas Ellenberg, PhD, Rui Feng, PhD, Melissa Fernando, MPH, Howard Panitch, MD,

Barbara Schmidt, MD, MSc Pamela Shaw, PhD, Ann Tierney, BA, MS

