## Supplemental Table and Supplemental Figure for "Lymphocyte-Specific Biomarkers Associated with Preterm Birth and Bronchopulmonary Dysplasia"

Gloria S. Pryhuber, MD

Division of Neonatology

University of Rochester Medical Center

Box 651, 601 Elmwood Avenue

Rochester, NY 14642, USA

***Supplemental Table 1:***

| **Definition** | **Criteria** |
| --- | --- |
| BPD (Shennan) | Supplemental oxygen requirement at 36 weeks PMA in infants born with birth weight (BW) < 1,500 g [1] |
| BPD (Physiologic) or Room Air Challenge (RAC) | Requirement of oxygen support (>21%) for at least 28 days and a subsequent assessment at 36 weeks PMA or discharge, whichever comes first. [2] |
| PRD | PRD was diagnosed if there were positive responses on at least 2 caregiver post-discharge questionnaires to the following: (1) hospitalization for respiratory indication, (2) home respiratory support, (3) respiratory medication administration, and/or (4) respiratory symptoms without a cold.[3] |
| Oxygen_AUC14_ | Calculated by the formula defined in Benaron and Benitz (1994) [4] using information recorded in the daily respiratory flowsheet data (FIO2, respiratory support mode, and applied airway pressure or cannula flow) through the first 28 days of life.[3] |

***Supplemental Table 1: Demographic and Diagnostic information for 130 subjects data from which have been used in transcriptomic analysis.***

| **Subject ID** | **Site** | **Gender** | **Hispanic Ethnicity** | **Race** | **Gest Age Birth (GAB) in Weeks** | **Shennan BPD** | **Physiologic BPD (RAC)** | **Oxygen_AUC14_**  **Oxygen Utilization at Day 14** | **PRD Status** |
| --- | --- | --- | --- | --- | --- | --- | --- | --- | --- |
| J9028 | Rochester | Male | No | White | 27.43 | No | No | 400.50 | Yes |
| J599B | Rochester | Male | No | White | 25.00 | Yes | No | 3204.00 | Yes |
| J061C | Rochester | Male | No | White | 28.57 | Unclassified | No | 22.00 | No |
| J84E4 | Rochester | Female | No | African American | 27.57 | No | No | 99.00 | Yes |
| JBD2A | Rochester | Female | No | African American | 27.57 | No | No | 39.00 | Yes |
| JAAD7 | Rochester | Female | No | White | 26.57 | Yes | Yes | 2358.50 | Yes |
| J94E8 | Rochester | Male | No | African American | 24.71 | No | Yes | 8124.50 | Yes |
| JD1FD | Rochester | Female | No | African American | 23.86 | No | No | 3887.00 | Yes |
| J96D6 | Rochester | Male | No | White | 25.43 | No | No | 1936.50 | Yes |
| J00F9 | Rochester | Female | No | White | 27.71 | No | No | 236.00 | No |
| J6B6F | Rochester | Male | No | African American | 26.71 | No | No | 2413.00 | Yes |
| J94F4 | Rochester | Female | No | African American | 26.00 | No | Yes | 1758.50 | No |
| JF22D | Rochester | Male | No | White | 27.14 | Yes | Yes | 3257.50 | Yes |
| J9977 | Rochester | Female | No | White | 27.71 | Unclassified | No | 340.50 | NA |
| J8DB7 | Rochester | Male | No | White | 30.43 | NA | NA | 0.00 | No |
| J2211 | Rochester | Male | No | Asian | 26.14 | No | No | 1072.00 | No |
| J835B | Rochester | Male | No | White | 25.57 | Yes | Yes | 1076.00 | Yes |
| J33AF | Rochester | Male | No | White | 25.57 | No | No | 1490.50 | No |
| J8CBE | Rochester | Female | No | White | 25.57 | No | No | 1344.00 | No |
| JA82D | Rochester | Female | No | White | 28.00 | Unclassified | No | 355.00 | Yes |
| J2B52 | Rochester | Female | No | Other Race | 30.14 | NA | NA | 188.53 | Yes |
| JB14F | Rochester | Female | No | White | 32.00 | NA | NA | 4.00 | Yes |
| J4D32 | Rochester | Male | No | White | 32.00 | NA | NA | 112.00 | Yes |
| J80AD | Rochester | Female | No | White | 28.86 | No | No | 333.50 | Yes |
| J8648 | Rochester | Female | No | White | 31.29 | NA | NA | 170.50 | Yes |
| J1B12 | Rochester | Female | No | African American | 24.14 | No | No | 2951.50 | No |
| J8C76 | Rochester | Female | Yes | White | 28.00 | No | No | 440.50 | No |
| JE573 | Rochester | Female | No | African American | 23.29 | No | No | 11204.50 | No |
| J6BE4 | Rochester | Male | No | White | 32.14 | NA | NA | 0.00 | No |
| JDCAF | Rochester | Female | No | White | 32.14 | NA | NA | 0.00 | No |
| JEACE | Rochester | Male | No | White | 32.14 | NA | NA | 664.00 | Yes |
| J1D4E | Rochester | Male | No | White | 24.14 | No | Yes | 3407.00 | Yes |
| J6213 | Rochester | Female | Yes | Other Race | 34.00 | NA | NA | 0.00 | No |
| J5633 | Rochester | Female | No | White | 32.14 | NA | NA | 22.50 | No |
| J92FE | Rochester | Female | No | African American | 35.14 | NA | NA | 37.00 | Yes |
| J5692 | Rochester | Male | No | African American | 31.57 | NA | NA | 339.00 | No |
| NEF69 | Rochester | Male | Yes | White | 34.71 | NA | NA | 0.00 | NA |
| ND6D4 | Rochester | Female | No | White | 33.29 | NA | NA | 0.00 | Yes |
| NE43D | Rochester | Male | No | White | 35.29 | NA | NA | 0.00 | Yes |
| J99F0 | Rochester | Female | No | White | 33.00 | NA | NA | 0.00 | No |
| J3150 | Rochester | Female | No | White | 33.57 | NA | NA | 0.00 | No |
| J0A30 | Rochester | Female | No | White | 32.86 | NA | NA | 0.00 | No |
| J8154 | Rochester | Male | No | White | 29.00 | NA | NA | 210.50 | Yes |
| N069F | Rochester | Male | Yes | Other Race | 35.00 | NA | NA | 1365.00 | No |
| E8990 | Rochester | Male | No | White | 39.14 | NA | NA | 0.00 | No |
| J1081 | Rochester | Male | No | White | 34.29 | NA | NA | 0.00 | NA |
| JEAC5 | Rochester | Female | No | White | 28.57 | Yes | Yes | 2552.50 | Yes |
| N52A3 | Rochester | Female | No | African American | 26.86 | No | No | 1080.00 | Yes |
| N174C | Rochester | Female | No | African American | 23.86 | Yes | Yes | 12501.50 | Yes |
| N05A1 | Rochester | Female | No | White | 26.14 | No | No | 1861.50 | No |
| ND064 | Rochester | Male | No | White | 40.29 | NA | NA | 0.00 | NA |
| JF17A | Rochester | Male | No | African American | 25.86 | No | No | 1452.50 | NA |
| NB0D2 | Rochester | Female | No | African American | 24.71 | No | No | 6826.00 | Yes |
| N2FF6 | Rochester | Male | No | White | 30.57 | NA | NA | 0.00 | No |
| NA9BF | Rochester | Male | No | White | 30.57 | NA | NA | 110.00 | NA |
| N85B5 | Rochester | Female | No | White | 30.57 | NA | NA | 179.00 | Yes |
| NAF88 | Rochester | Female | No | White | 39.29 | NA | NA | 0.00 | No |
| NA54B | Buffalo | Female | No | White | 31.29 | NA | NA | 0.00 | Yes |
| NF46C | Buffalo | Male | No | White | 25.86 | Yes | Yes | 8483.00 | Yes |
| N0E97 | Buffalo | Female | No | White | 26.57 | Yes | Yes | 6067.00 | Yes |
| N8C7D | Buffalo | Male | No | White | 31.43 | NA | NA | 432.92 | Yes |
| N4696 | Buffalo | Male | Yes | Other Race | 28.00 | Yes | Unclassified | 7650.50 | No |
| ND11C | Buffalo | Male | No | Other Race | 24.71 | Yes | No | 17486.50 | NA |
| N8248 | Buffalo | Female | No | White | 25.43 | Yes | No | 6722.50 | Yes |
| ND69A | Buffalo | Male | No | African American | 25.43 | No | No | 8038.50 | Yes |
| N9B2B | Buffalo | Male | No | White | 24.57 | No | Yes | 9628.00 | Yes |
| NFBC1 | Buffalo | Male | No | White | 28.71 | Unclassified | No | 1019.50 | Yes |
| N09F5 | Buffalo | Male | No | White | 28.71 | Yes | Yes | 3199.00 | Yes |
| N5FCD | Buffalo | Male | No | Asian | 28.57 | No | No | 775.00 | No |
| N2F4C | Buffalo | Male | No | Asian | 28.57 | No | No | 385.50 | Yes |
| N0987 | Buffalo | Male | No | White | 31.71 | NA | NA | 0.00 | Yes |
| N840F | Buffalo | Female | No | African American | 25.00 | Yes | Yes | 7025.50 | No |
| NDBE8 | Buffalo | Male | No | White | 32.71 | NA | NA | 120.14 | Yes |
| N6ECA | Buffalo | Male | No | White | 32.71 | NA | NA | 0.00 | Yes |
| NBE6F | Buffalo | Female | No | White | 28.14 | Yes | Yes | 618.50 | Yes |
| ND48A | Buffalo | Female | No | White | 27.43 | No | Yes | 304.00 | Yes |
| ND0F8 | Buffalo | Female | No | White | 27.43 | No | No | 1616.50 | No |
| NEA48 | Buffalo | Male | No | White | 32.14 | NA | NA | 0.00 | NA |
| N8220 | Buffalo | Male | No | African American | 28.14 | Unclassified | No | 869.50 | No |
| N81A1 | Buffalo | Male | No | African American | 28.14 | Unclassified | No | 1831.46 | No |
| ND834 | Buffalo | Female | No | White | 28.71 | Unclassified | No | 0.00 | Yes |
| N7E07 | Buffalo | Female | No | White | 28.43 | Yes | Yes | 4139.50 | Yes |
| N6A6C | Buffalo | Female | No | White | 26.14 | Yes | No | 2799.00 | No |
| NB35B | Buffalo | Male | No | White | 26.86 | No | No | 123.00 | Yes |
| NF65F | Buffalo | Male | No | White | 25.43 | Yes | Yes | 3853.00 | No |
| N9ACD | Buffalo | Male | No | African American | 26.14 | Yes | Yes | 8016.00 | Yes |
| N5688 | Buffalo | Female | No | African American | 26.14 | Yes | Yes | 7685.00 | Yes |
| NEB8C | Buffalo | Male | No | White | 31.86 | NA | NA | 0.00 | No |
| NB109 | Buffalo | Male | No | African American | 24.71 | No | No | 9626.00 | Yes |
| NC6F1 | Buffalo | Male | No | White | 34.29 | NA | NA | 0.00 | NA |
| NEF4C | Buffalo | Male | No | African American | 34.86 | NA | NA | 0.00 | No |
| NCCDE | Buffalo | Female | No | White | 28.43 | No | No | 1735.81 | No |
| NC52C | Buffalo | Female | No | African American | 29.29 | NA | NA | 0.00 | Yes |
| N477D | Buffalo | Female | No | African American | 27.14 | No | Yes | 10201.00 | Yes |
| NA845 | Buffalo | Female | No | White | 27.00 | Yes | Yes | 2656.00 | No |
| N0C8F | Buffalo | Male | No | White | 28.00 | Yes | Yes | 6635.50 | Yes |
| N83DA | Buffalo | Female | No | White | 33.86 | NA | NA | 0.00 | NA |
| N679A | Buffalo | Male | No | African American | 26.57 | Yes | Unclassified | 2406.00 | Yes |
| NC8D9 | Buffalo | Female | No | White | 28.14 | Yes | Unclassified | 2620.00 | Yes |
| N3037 | Buffalo | Male | No | White | 25.14 | Yes | No | 9185.50 | No |
| N51DF | Buffalo | Male | No | White | 33.86 | NA | NA | 67.50 | Yes |
| NFFFF | Buffalo | Male | No | African American | 35.71 | NA | NA | 0.00 | No |
| NB735 | Buffalo | Female | No | White | 33.14 | NA | NA | 851.50 | No |
| ND4BE | Buffalo | Male | No | White | 33.14 | NA | NA | 1369.50 | No |
| NC2D3 | Buffalo | Female | No | White | 31.00 | NA | NA | 123.50 | Yes |
| N9932 | Buffalo | Female | No | African American | 26.14 | No | No | 5172.00 | Yes |
| NDE39 | Buffalo | Female | No | White | 32.71 | NA | NA | 0.00 | No |
| N880B | Buffalo | Female | No | African American | 32.00 | NA | NA | 0.00 | Yes |
| N7D89 | Buffalo | Female | No | African American | 32.00 | NA | NA | 0.00 | Yes |
| N2001 | Buffalo | Male | No | African American | 27.71 | No | No | 472.00 | Yes |
| N2C5E | Buffalo | Female | No | White | 28.71 | Unclassified | Unclassified | 421.12 | Yes |
| N7764 | Buffalo | Female | Yes | Other Race | 24.00 | Yes | Yes | 12359.50 | Yes |
| N70E9 | Buffalo | Male | No | White | 26.00 | No | No | 3743.50 | Yes |
| NB272 | Buffalo | Female | Yes | Other Race | 32.86 | NA | NA | 0.00 | NA |
| NEC86 | Buffalo | Female | Yes | Other Race | 32.86 | NA | NA | 0.00 | NA |
| N046E | Buffalo | Male | No | White | 25.14 | Yes | Yes | 2379.00 | Yes |
| N7982 | Buffalo | Male | No | White | 25.14 | No | No | 3239.50 | No |
| NB903 | Buffalo | Female | No | African American | 25.43 | No | Yes | 8133.00 | Yes |
| N003F | Buffalo | Male | No | African American | 25.43 | Yes | Yes | 13007.50 | Yes |
| N1CE2 | Buffalo | Female | No | African American | 25.43 | Yes | Yes | 6948.00 | Yes |
| N6703 | Buffalo | Male | No | African American | 34.43 | NA | NA | 0.00 | No |
| ND977 | Buffalo | Male | No | African American | 35.43 | NA | NA | 0.00 | NA |
| NB4EE | Buffalo | Male | No | White | 41.29 | NA | NA | 0.00 | No |
| N7F32 | Buffalo | Male | No | White | 41.14 | NA | NA | 0.00 | Yes |
| N49C4 | Buffalo | Female | No | White | 41.14 | NA | NA | 0.00 | No |
| N720A | Buffalo | Female | No | White | 30.86 | NA | NA | 44.50 | Yes |
| N854C | Buffalo | Female | No | White | 40.43 | NA | NA | 0.00 | No |
| NBF2B | Buffalo | Male | No | African American | 34.00 | NA | NA | 0.00 | Yes |
| N2749 | Buffalo | Male | No | African American | 40.57 | NA | NA | 0.00 | No |
| N2895 | Buffalo | Male | No | White | 40.14 | NA | NA | 0.00 | No |

***Supplemental Figure 1: The steps involved in the study and the applications used.***

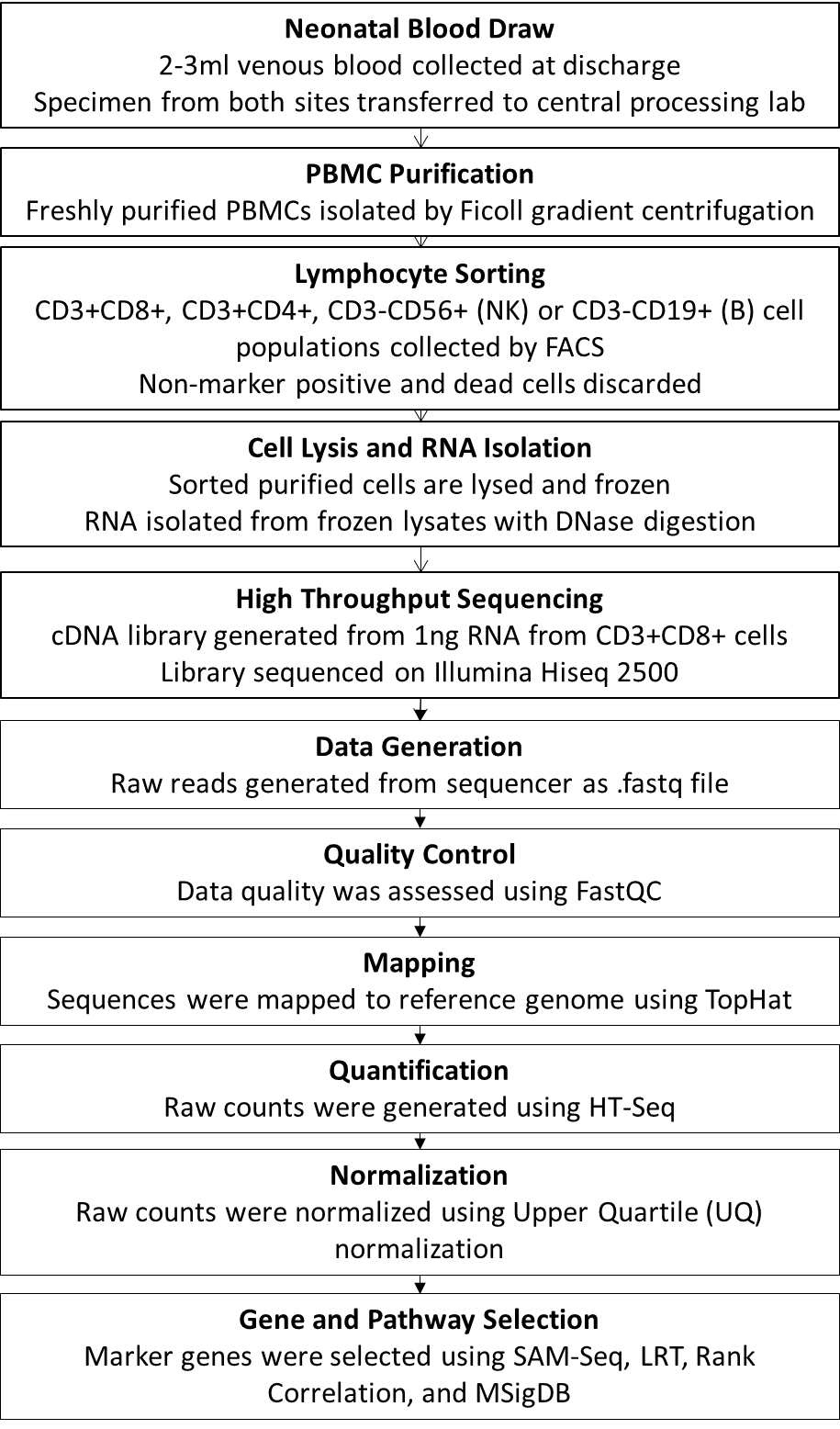

***Supplemental Figure 2: RNA-Seq quality statistics: High throughput sequencing was performed using RNA isolated from sorted CD8 + T cells (N=145). The number of input reads (A), the rate of reads mapped (B) and proportion of the genome represented by expressed transcripts (C) for each sample is displayed.***

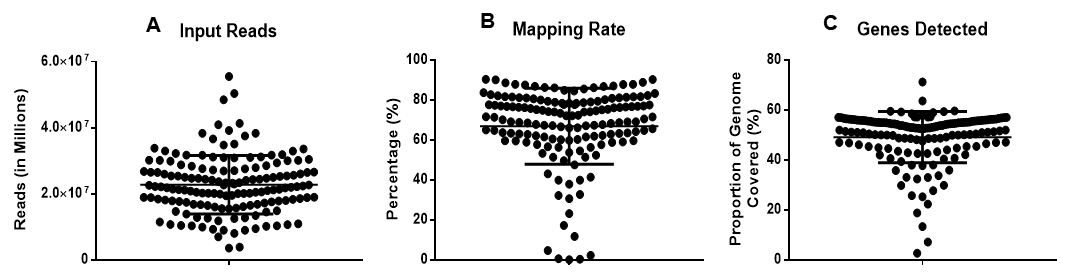

***Supplemental Figure 3: Predictive markers for BPD. A) Normalized expression for 92 genes is shown with subjects in individual columns and genes in individual rows. High expression is indicated by red color and low expression is indicated by green. B) Plots of area under the receiver-operator curve (AUC) comparing the performance of three predictors; using subject gestational age (GA) only, using pathway-based gene expression selected by screened PCA (sPCA) only, or using a combination of GA and sPCA data.***

A

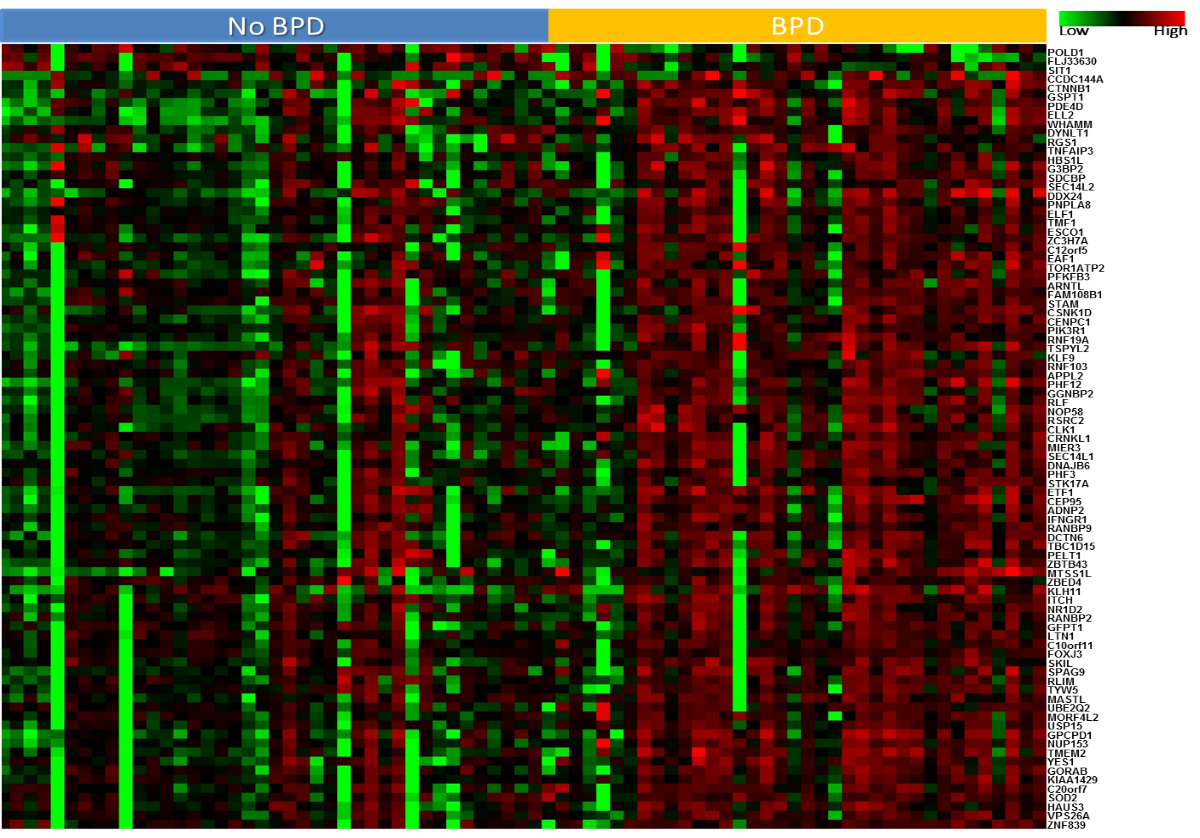

B

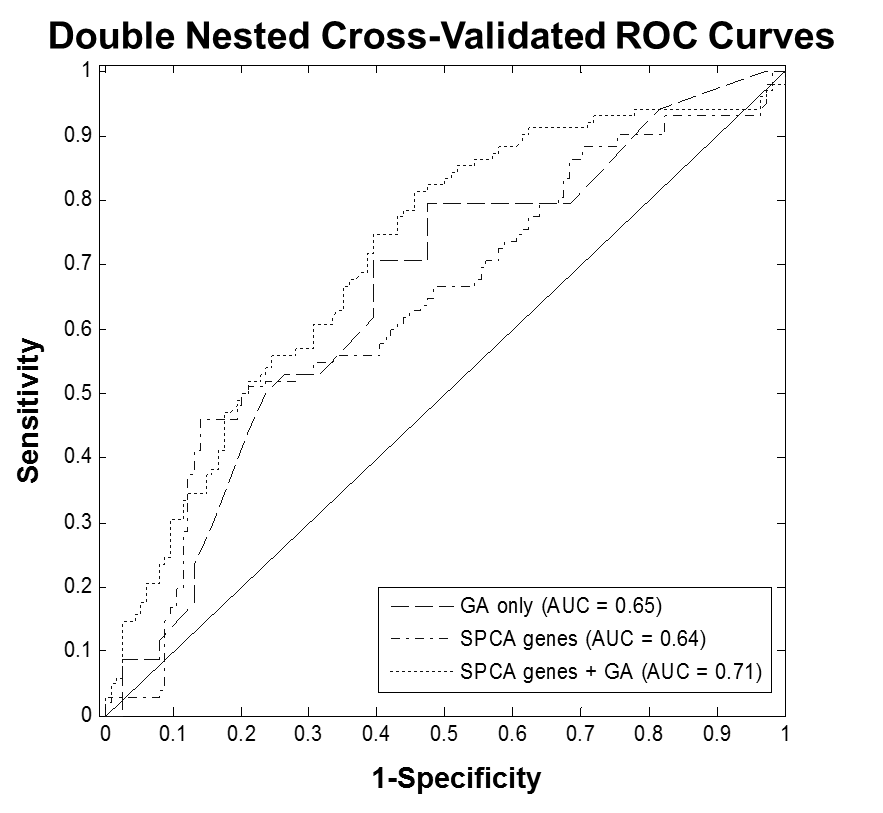

***Supplemental Figure 4: Distribution of number of genes differentially expressed in BPD, as assessed by the multiple analytical approaches using the different clinical and physiological definitions of BPD in subjects less than 29 weeks of gestation at the time of birth. O2 (Correlation) indicates correlation analysis with oxygen utilization, RAC (SAM) indicates differentially expressed genes identified by SAM-Seq in BPD as defined by RAC, Shennan (SAM) indicates differentially expressed genes identified by SAM-Seq in BPD as defined by the Shennan definition, and BPD-GAB (sPCA) indicates genes identified as differentially expressed by sPCA in BPD after age adjustment. Irrespective of the method used, or the clinical or physiological definition applied, nine genes were consistently different in BPD subjects.***

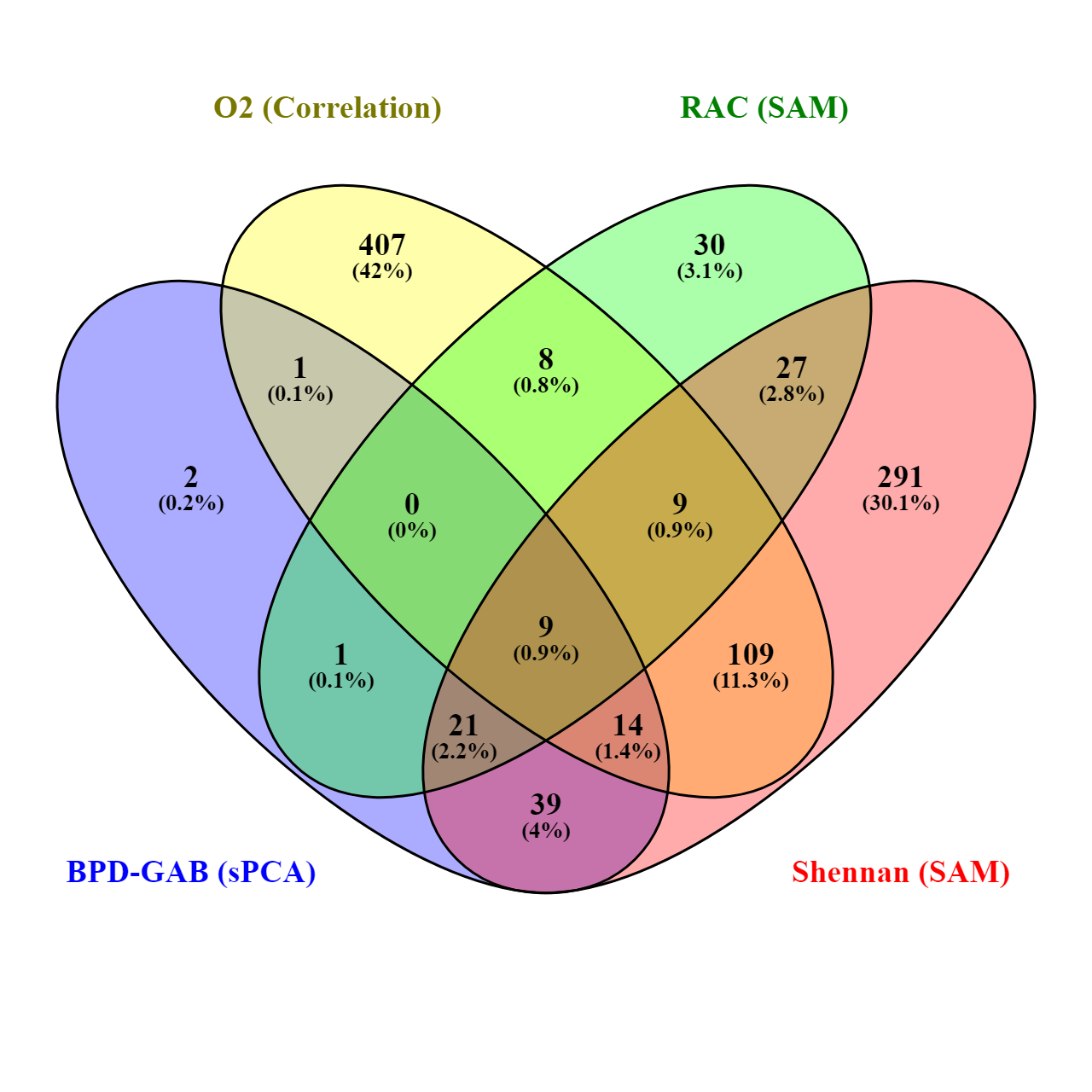
